# A NOVEL DEEP LEARNING MODEL, RDB CYCYLEGAN-CBAM FOR LOW-DOSE CT IMAGE DENOISING

**DOI:** 10.64898/2026.02.17.706311

**Authors:** Osman Assaf, Albert Güveniş

## Abstract

Computed Tomography (CT) is one of the largest contributors to radiation exposure from medical imaging, which can induce DNA damage and increase cancer risk. Reducing CT radiation dose to improve patient safety inherently increases image noise and artifacts. Generative adversarial networks (GANs) have shown promise for unsupervised low-dose CT (LDCT) denoising. Building on this, RDBCycleGAN-CBAM, a CycleGAN-based model that integrates residual dense blocks (RDBs) and convolutional block attention modules (CBAM), was developed to effectively denoise quarter-dose CT images while preserving structural detail. The model was trained on unpaired quarter-dose and full-dose CT scans from the NIH-AAPM-Mayo dataset using adversarial (LSGAN), cycle-consistency, and identity losses. Evaluation on held-out test slices was performed using PSNR and SSIM as the primary image-quality metrics. The results demonstrate that the proposed RDBCycleGAN-CBAM method not only achieves higher peak signal-to-noise ratio (PSNR) and structural similarity index (SSIM) values but also outperforms most existing deep learning-based methods, achieving mean improvements of +3.97 dB in PSNR and +0.053 in SSIM relative to quarter-dose inputs. Shapiro– Wilk tests for PSNR and SSIM motivated the use of the nonparametric Wilcoxon signed-rank test, by which highly significant improvements across both metrics (PSNR and SSIM) were demonstrated. The very large rank-biserial correlation values (1.0) indicate that nearly all test images experienced substantial quality improvement. Furthermore, the narrow bootstrap confidence intervals for the mean differences suggest that these improvements are consistent across the dataset. These advancements contribute to medical imaging by providing a viable, vendor-neutral tool for reducing patient radiation exposure without compromising diagnostic value.

## I. INTRODUCTION

Computed tomography (CT) has become an indispensable diagnostic imaging tool in modern medicine, widely used for rapid, detailed assessment of diverse conditions [1]. Despite comprising only a fraction of imaging procedures, CT accounts for a disproportionate share of patient radiation exposure [2], [3]. Consequently, guidelines emphasize the “as low as reasonably achievable” (ALARA) principle, seeking to minimize dose, especially in sensitive populations, while preserving diagnostic utility [3]. Lowering CT dose directly reduces X-ray photon counts, which inevitably degrades the image signal-to-noise ratio and introduces reconstruction artifacts [4]. Indeed, low-dose CT (LDCT) images are known to contain substantially increased noise and artifacts compared to normal-dose scans, which can obscure lesions and reduce diagnostic confidence [5]. Hence, there is a critical need for advanced image processing techniques that can suppress noise and artifacts in LDCT data while preserving the underlying anatomical and pathological detail [6],[7]. Over the past decade, a plethora of denoising methods have been developed for LDCT, ranging from iterative reconstruction and filtering approaches to machine learning models [8]. In particular, generative adversarial networks with cycle consistency (CycleGAN) have proven effective, they learn a bidirectional mapping between low-dose and high-quality CT domains using unpaired datasets [9],[10]. Despite this progress, prior CycleGAN-based denoisers have notable drawbacks. CycleGAN often produces oversmoothed results, where fine anatomical details and texture can be lost or blurred [10], [11]. Moreover, many CycleGAN denoisers lack explicit mechanisms to focus on clinically important structures, risking artifacts or loss of diagnostic content [11]. To address the shortcomings of prior methods, we propose RDBCycleGAN-CBAM: a novel cycle-consistent GAN tailored for LDCT denoising. Our generator networks incorporate Residual-Dense Blocks (RDBs)—densely connected convolutional layers with local feature fusion—which have been shown to extract rich hierarchical features and stabilize training in image super-resolution tasks [12]. We also embed a Convolutional Block Attention Module (CBAM) in the network bottleneck, which applies sequential channel-wise and spatial attention to refine feature maps [13]. Additionally, we include dilated convolutions in the mid-level layers to enlarge the receptive field without additional pooling, helping the network capture broader contextual information [14]. Compared to a vanilla CycleGAN, our model has a more compact generator (fewer parameters via the shared RDB structure) and a focused attention module, which together promote stable, detail-preserving denoising.

## II. METHODOLOGY

The proposed RDBCycleGAN-CBAM framework follows an unpaired CycleGAN paradigm for low-dose CT denoising. We employ two generator networks, G_Q→F_ (quarter-dose to full-dose) and G_F→Q_ (full-dose to quarter-dose) as shown in Fig. 1, each mirroring the other architecture. Each generator is based on an encoder–decoder with dense skip connections and a specialized bottleneck. In the encoder and decoder stages we stack Residual-Dense Blocks (RDBs), which are inspired by the Residual Dense Blocks of RDN but pruned for computational efficiency. Each RDB connects all its convolutional layers densely and adds a local residual connection, enabling contiguous memory of features [16]. These dense connections allow the network to exploit hierarchical features from all layers, enhancing feature reuse and gradient flow [16],[17]. Between the encoder and decoder, we insert a dilated convolutional block with dilation rates >1 to expand the receptive field without loss of resolution [14]. The decoder mirrors the encoder with upsampling layers (nearest-neighbor or transposed convolutions) and skip connections from encoder layers to corresponding decoder layers [16].

**Fig. 1.**
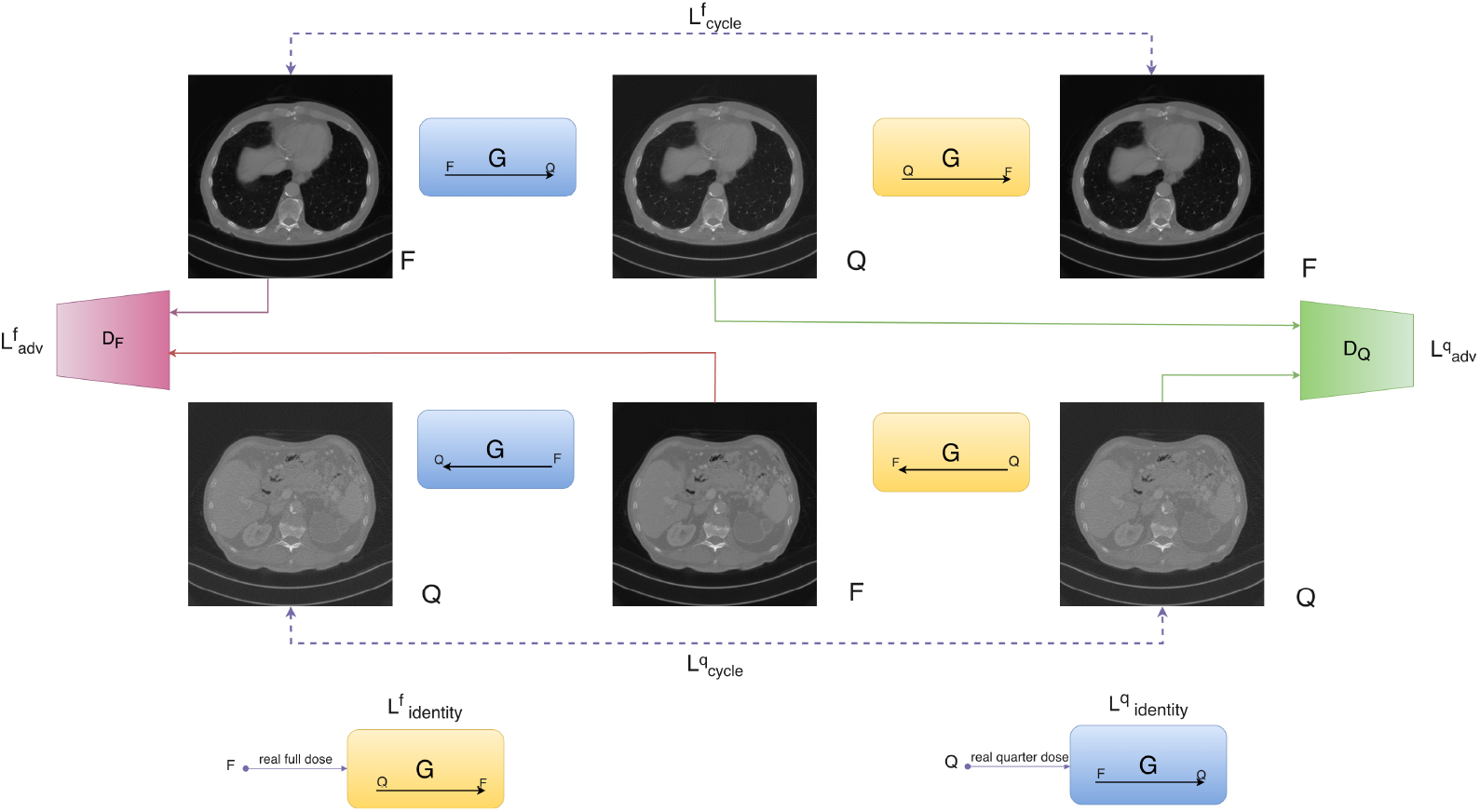
The overall architecture CycleGAN network for LDCT image denoising. There are two generators G_F→Q_ and G_Q→F_, their corresponding discriminators, D_Q_, D_F_, anft he loss functions L_cycle_, L_adv_ and L_identity_

To further improve feature representation, we integrate the Convolutional Block Attention Module (CBAM)[18] within each generator[9]. CBAM sequentially computes spatial and channel-wise attention maps on feature maps, refining salient features with negligible overhead[14]. Specifically, after each RDB sequence and before upsampling, a CBAM refines the feature tensor by first applying channel attention (global average/max pooling followed by shared MLP) and then spatial attention (convolution over aggregated channel information). This attention mechanism helps the network emphasize important structures (e.g. edges, textures) and suppress irrelevant noise, aligning with prior findings that CBAM can improve representation in diverse vision tasks [9].

### A. Generators

The model uses two symmetric generators, G_F→Q_ and G_Q→F_, which map between the full-dose (F) and quarter-dose (Q) image domains. Each generator is an encoder–decoder network with A bottleneck designed and integrated in the deepest level of the generator. Fig. 4, A single input channel is mapped to a single output channel. The base channel width g_channels_ = 32 is multiplied by factors [1, 2, 4, 8] at successive resolutions.

#### 1. Encoder

The encoder begins with a 3 × 3 convolution (stride 1) mapping the input to 32 channels. It then proceeds through four resolution levels. Fig. 2, At each level, a sequence of ResNet blocks (each consisting of two 3 × 3 convolutions with GroupNorm and ReLU, plus a 1 × 1 convolution shortcut) is applied. After the ResNet blocks at levels 1–3, a downsampling convolution (3×3, stride 2) is applied to halve the spatial dimensions. The channel dimension is multiplied by the levelâs factor (1 → 2 → 4 → 8). This multi-resolution design allows the model to capture features at different scales.

**Fig. 2.**
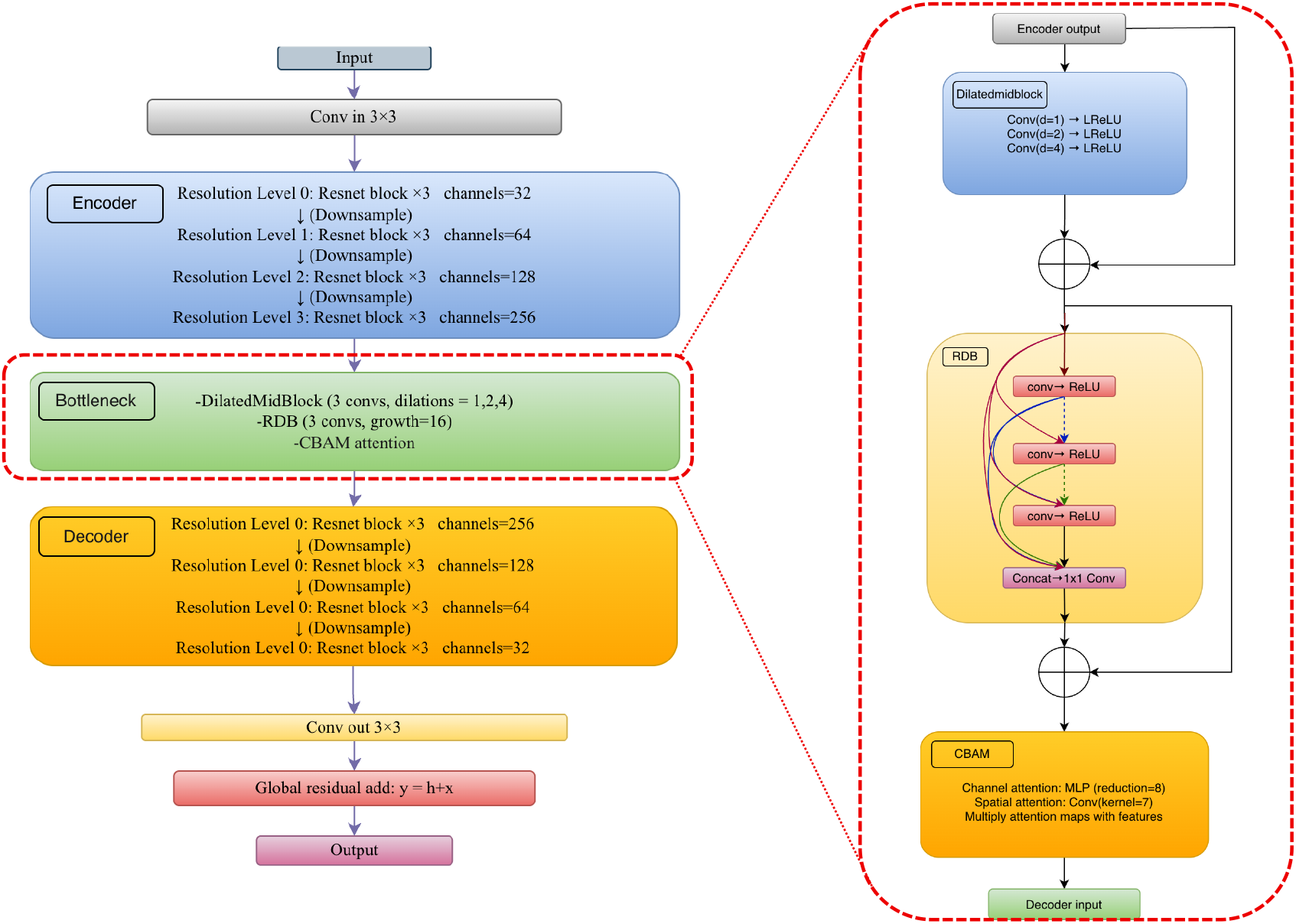
The proposed Generator Architecture.

#### 2. Bottleneck

At the coarsest scale (channel multiplier 8), the encoder output passes through a DilatedMidBlock and a LightRDB, followed by an optional CBAM attention module. The DilatedMidBlock consists of three 3×3 convolutions with dilation factors 1, 2, and 4 (padding equal to dilation) to enlarge the receptive field without further downsampling. This is followed by a lightweight Residual Dense Block (LightRDB) with three densely connected 3×3 convolutional layers (growth rate 16) and a 1×1 convolution to restore the channel dimension. The LightRDB’s output is added (scaled by 0.2) to its input as a residual. Finally, a Convolutional Block Attention Module (CBAM) applies channel-wise and spatial attention to refine the bottleneck features as shown in Fig. 3.

**Fig. 3.**
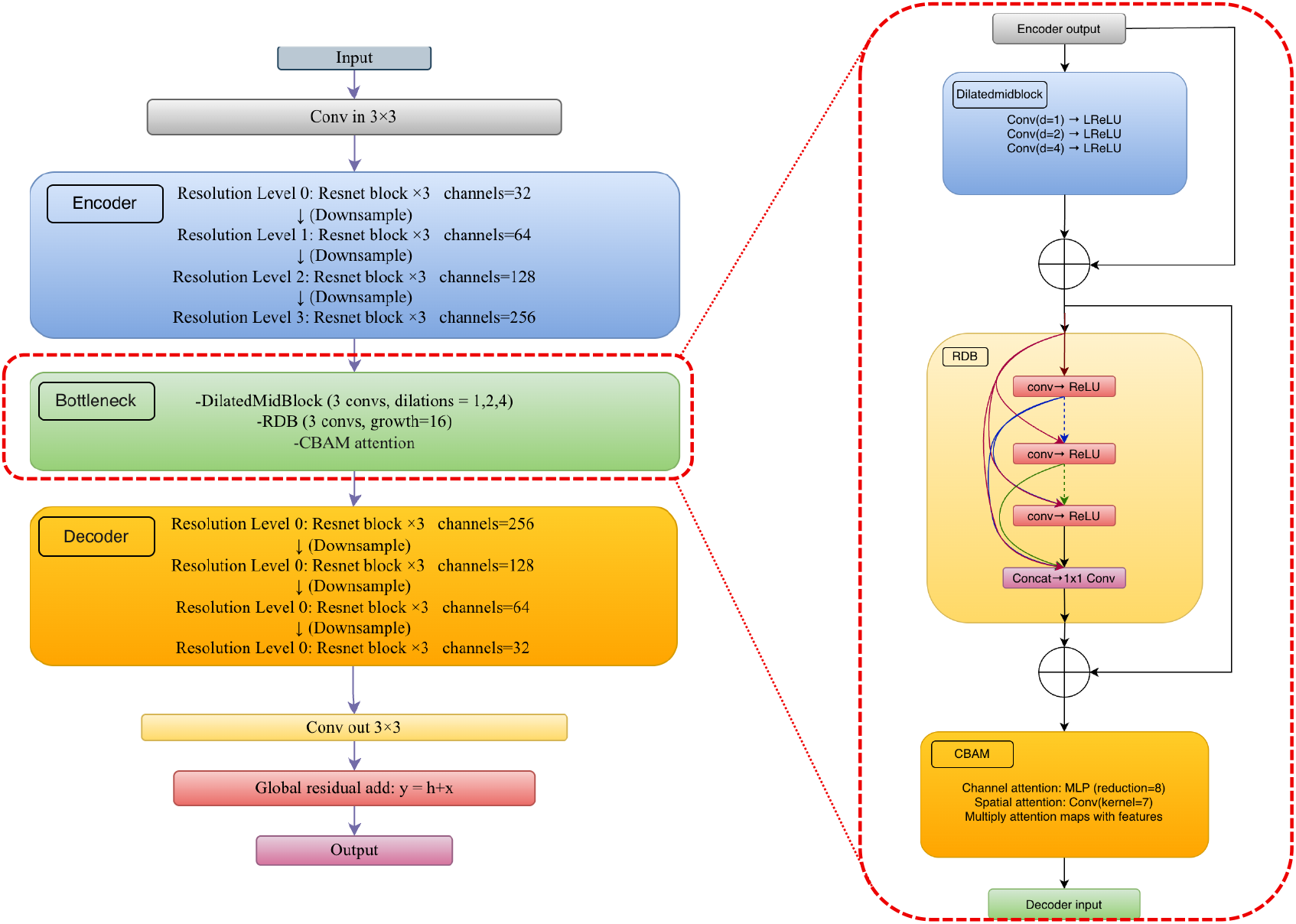
Proposed bottleneck architecture integrated at the deepest stage of the generator.

#### 3. Decoder

The decoder mirrors the encoder. At each level (from coarse to fine), we concatenate the decoders current feature map with the corresponding encoder feature map (skip connection) and pass the concatenated tensor through ResNet blocks. When moving to a finer scale, we apply bilinear upsampling: specifically, each upsampling block consists of a 2× bilinear upsampling layer followed by a 3 × 3 convolution, batch normalization, and LeakyReLU(0.1). The channel dimensions are reduced by the same factors (8→4→2→1) in reverse [11].

#### 4. Global Residual Skip

After the final upsampling, a 3 × 3 output convolution maps the features back to one channel. The generator uses a global residual connection: the network’s output h is added to the original input x to form the final output y = h+x, Fig. 2. This helps the network focus on learning the noise to add/subtract rather than duplicating the entire image content.

### B. Discriminators

For the discriminators D_Q_, D_F_, we adopt a PatchGAN architecture [15]. Each discriminator is a convolutional network that classifies overlapping image patches (e.g. 70 × 70 pixel regions) as real or fake. Our discriminator has five layers, Fig. 5, it gradually downsamples the input (low-dose or full-dose image) by factors of 2 using 4 × 4 convolutional kernels with stride 2 (except the last layers use stride 1) and uses Instance Normalization in intermediate layers. This results in an N × N grid of output probabilities rather than a single score.

**Fig. 4.**
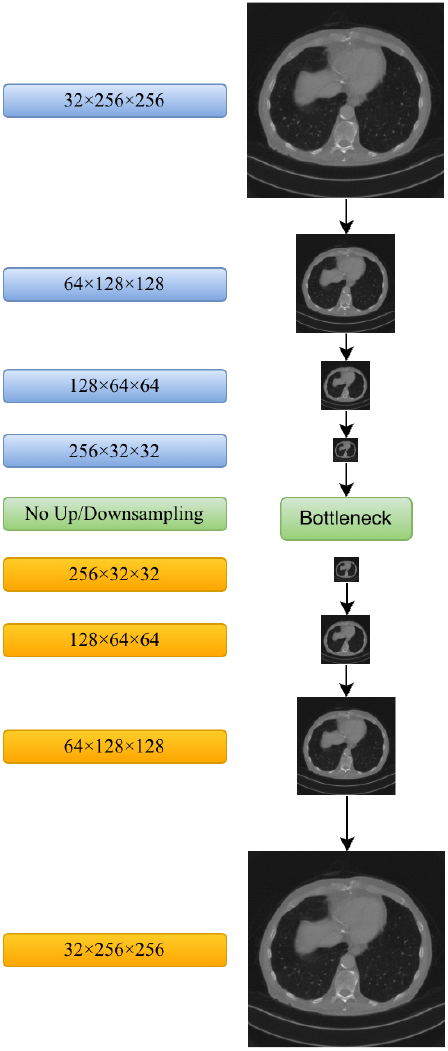
Image Forward pass Proceeding in the Generator.

**Fig. 5.**
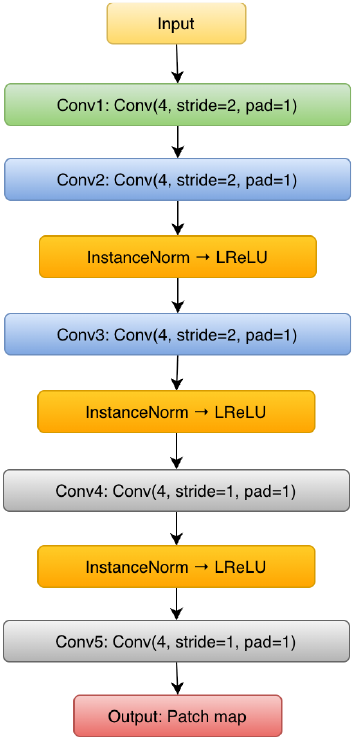
The Architecture of the Discriminator.

PatchGAN effectively models high-frequency differences; in our case it helps ensure the output CT patches have realistic noise texture and edges at the local level. We set the base number of discriminator filters to 64, doubling at each layer (64→128→256→512) as is standard. Leaky ReLU is used as the activation. By restricting its focus to patches, the discriminator guides the generator to produce outputs that are plausible in a local sense, which is crucial for medical image fidelity.

### C. LOSS FUNCTIONS

The network is trained adversarially using a CycleGAN loss scheme [19]. The adversarial loss for each generator– discriminator pair uses the Least-Squares GAN (LSGAN) formulation [12] for stable training. Concretely, for generator GQ2F and its discriminator DF, the LSGAN loss is:

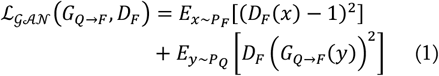

where x denotes a real full-dose CT image, y denotes a quarter-dose CT image, and DF (·) outputs patch-level realism scores. A similar adversarial loss, LGAN(G_F→Q_, DQ), is applied for the reverse translation direction. The least-squares GAN (LSGAN) formulation replaces the original binary cross-entropy loss with a squared error loss, which mitigates vanishing gradients and promotes more stable training convergence[20]. Each generator seeks to minimize its adversarial loss against its corresponding discriminator, while each discriminator aims to maximize this objective by assigning a target value of 1 to real images and 0 to generated (fake) images.

To ensure that the learned mappings are cycle-consistent, a cycle-consistency loss is incorporated. Specifically, for an input quarter-dose image y, the forward mapping G_Q→F_ (y) should be able to reconstruct the original image when subsequently passed through the reverse mapping G_F→Q_. This constraint enforces structural consistency and prevents arbitrary mappings between the two domains. Formally, the cycle-consistency loss is defined as[19]:

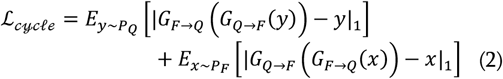

This uses the L1 norm (mean absolute error) to encourage reconstructed images to be close to the originals. Empirically, cycle consistency ensures that the two generators are inverses of each other on the data manifold, which is critical in the unpaired setting[20].

We also include an identity loss to encourage the generators not to alter images that are already in the target domain. That is, feeding a full-dose image x into G_Q→F_ (which should map quarter-dose to full-dose) should ideally return x unchanged, and similarly G_F→Q_ (y) ≈ y for quarter-dose y. Formally:

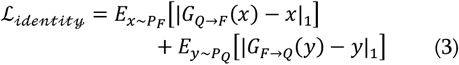

As noted in prior work, identity loss helps preserve the color/intensity calibration and anatomical detail when translating between medical image domains. It also stabilizes training by penalizing unnecessary deviations.

The total generator loss is a weighted sum of these components:

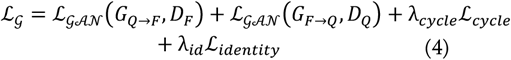

where λ_cycle_ and λ_id_ are weighting hyperparameters that balance the contributions of the cycle-consistency and identity losses, respectively. Following common practice in the literature, these were set to λ_cycle_ = 10 and λ_id_ = 5 to strongly enforce cycle consistency while preserving intensity fidelity. The discriminator objectives correspond to the standard least-squares GAN (LSGAN) discriminator losses associated with LGAN.

### D. Evaluation Metrics

#### 1. Peak Signal-to-Noise Ratio (PSNR)

PSNR measures peak image fidelity. For two images Itest and Iref with pixel values in [0, 1], PSNR is defined in terms of the mean squared error (MSE) as [21]:

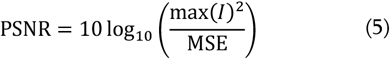

where max(I) is the maximum possible pixel value (here 1) and

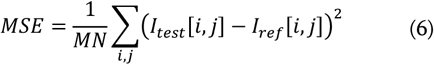

#### 2. Structural Similarity Index (SSIM)

SSIM measures perceptual similarity between two images by combining luminance, contrast, and structural comparison. For image patches x and y with means μ_x_,μ_y_, variances σ_x_^2^,σ_y_^2^, and covariance σ_xy_, SSIM is defined as [21]:

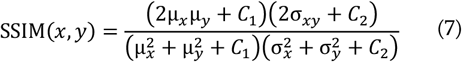

where C1 and C2 are small stabilizing constants[54]. SSIM ranges from −1 to 1, with higher values indicating greater similarity.

#### 3. Root-Mean-Square Error (RMSE) and Mean Absolute Error (MAE)

RMSE and MAE are standard pointwise error metrics. For two images a and b consisting of N pixels [22],

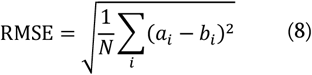

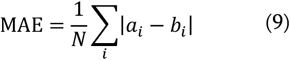

Both metrics quantify absolute reconstruction error, expressed here in Hounsfield units (HU).

#### 4. Normalized Mean Squared Error (NMSE)

NMSE scales the mean squared error relative to the energy of the reference image [23]. For images a and b,

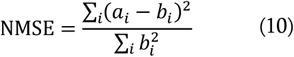

Lower NMSE indicates greater similarity, with NMSE = 0 only when the two images are identical.

#### 5. Signal-to-Noise Ratio (SNR)

For a reference image I_ref_ and a denoised image I_test_, the SNR (in dB) is defined as [21]:

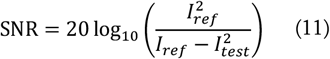

where ·2 denotes the Euclidean (2) norm over all pixel values. A larger SNR indicates a stronger signal relative to noise.

#### 6. Edge-Keeping Index (EKI)

EKI quantifies improvement in edge preservation. Let EdgeErr_in_ and EdgeErr_out_ denote the edge errors of the input and output images, respectively, computed as the L_1_ difference between gradient magnitudes (e.g., using Sobel filters) relative to the reference image [24]. The EKI is defined as:

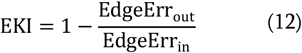

By this definition, the input baseline satisfies EKI = 0, while values closer to 1 indicate substantially better edge preservation in the output than in the input.

## III. IMPLEMENTATION

### A. Data Preparation

We trained and evaluated our model on the NIH-AAPM-Mayo Clinic Low Dose CT Challenge dataset [25]. This dataset provides 2D CT slice images of the human abdomen at normal dose and at a quarter of that dose. The dataset comprises 4,260 slices from ten anonymized patients with a slice thickness of 1.0, mm. We used 3,839 (8 patients) slice pairs for training and 421 pairs (2 patients) for testing (these were split by patient scans, so that test patients were not seen during training). All images are contrast-enhanced and were acquired during the portal venous phase using a Siemens SOMATOM Flash scanner. The images were originally reconstructed at 512 × 512 resolution with 1 mm slice thickness. Each CT slice is resized/cropped to 256 × 256 pixels. We normalize intensities to [0, 1] by clipping to the standard Hounsfield unit window and dividing by the maximum.

Pixel intensities were converted to Hounsfield Units and normalized for stable network training. Identical preprocessing was applied to both domains.

### B. Experimental Settings

The training conducted using a single high-performance GPU (NVIDIA Tesla A100, 32 GB VRAM), with a host system containing 12 GB system RAM and 256 GB SSD storage for datasets and checkpoints. PyTorch (v1.x) served as the deep learning framework. Under this configuration, training the RDBCycleGAN–CBAM model required approximately 22 hours. Mixed-precision training improved computational speed and reduced GPU memory usage without degrading numerical accuracy.

### C. Training Procedure

We wrote a training loop that iteratively fetched batches from the Data Loader, performed forward passes for all four network components (2 generators, 2 discriminators), computed the respective losses, and updated parameters. We used gradient penalty or normalization techniques as needed to keep the training stable (though in practice, the LSGAN + cycle consistency was stable). PyTorch’s automatic differentiation was utilized to backpropagate the weighted sum of losses. We also employed gradient scaling (AMP) to prevent floating-point underflow with mixed precision, Fig.7. During training, we periodically saved model checkpoints and logged losses.

**Fig. 6.**
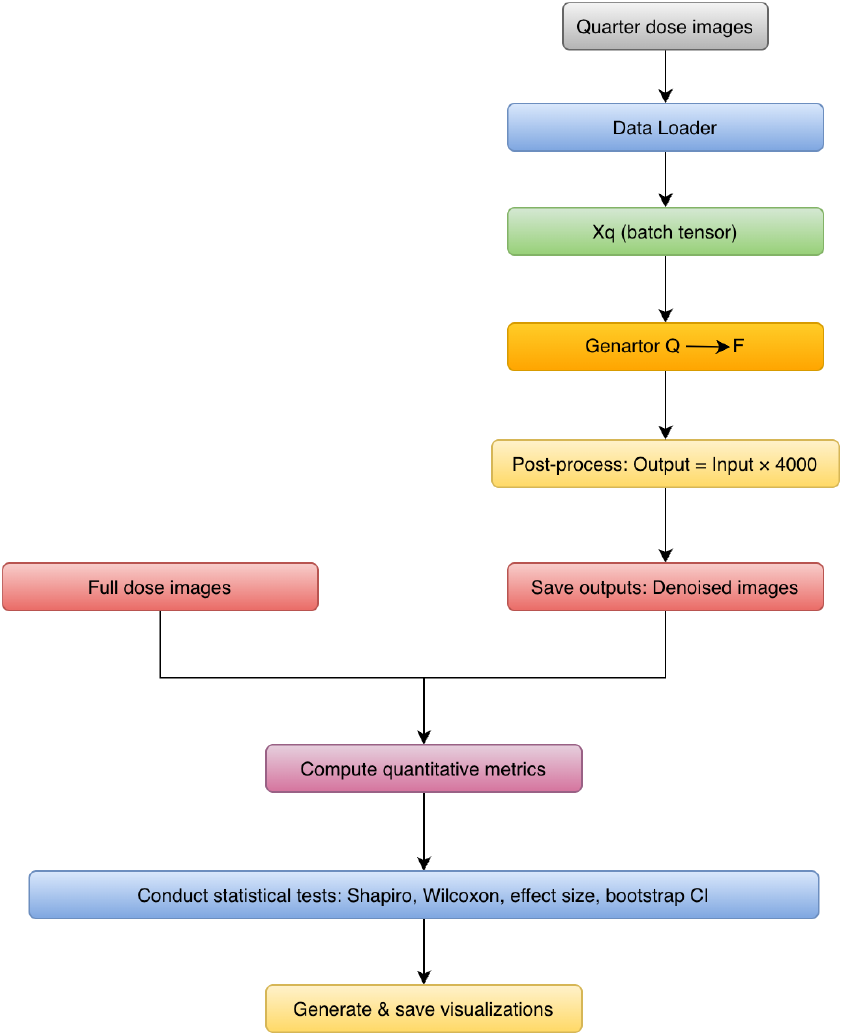
Training Workflow of the RDBCycleGAN-CBAM Model.

**Fig. 7.**
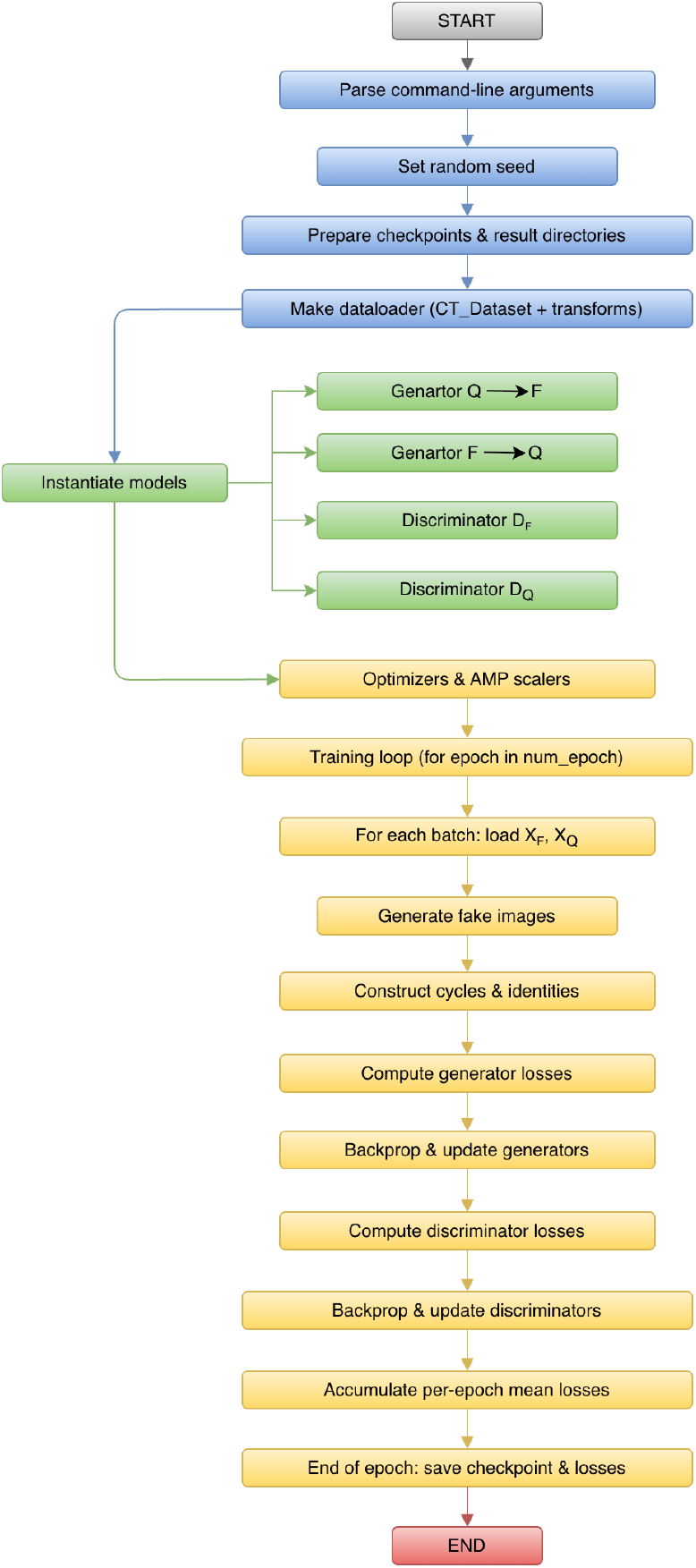
Training Workflow of the RDBCycleGAN-CBAM Model.

### D. Testing Procedure

The test pipeline loads the trained quarter→full generator (G_Q→F_) from a CycleGAN framework and performs slice-wise inference on low-dose CT inputs. The generator an encoder– bottleneck–decoder residual network (with dilated and residual dense blocks and attention) learned during training produces denoised, single channel outputs which are rescaled to HU-like values and stored per slice, Fig. 6. The script computes a comprehensive set of quantitative metrics (PSNR, SSIM, RMSE, MAE, NMSE, SNR, Sobel-based edge-error, and an edge-keeping index), conducts paired statistical tests and bootstrap confidence intervals, and generates multiple visualizations (residual maps, qualitative panels, Bland– Altman plots, and paired-lines plots) to summarize image quality improvements over the low-dose input.

## IV. RESULTS

### A. Quantitative Evalotion

Table II summarizes the quantitative comparison among LDCT inputs, vanilla CycleGAN, and the proposed RDBCycleGAN-CBAM. While vanilla CycleGAN improves image quality relative to LDCT across all metrics, the proposed method consistently achieves the best performance. Specifically, RDBCycleGAN-CBAM attains the highest PSNR (38.14 ± 4.54 dB) and SSIM (0.946 ± 0.033), along with the lowest RMSE (40.29 ± 8.58 HU), MAE (30.03 ± 6.85 HU), and NMSE (0.003 ± 0.002). It also yields the highest SNR and the lowest edge error, with an Edge-Keeping Index (EKI) of 0.428, indicating superior noise suppression while preserving structural detail. Overall, the results demonstrate that the integration of residual-dense blocks and attention mechanisms provides consistent and substantial improvements over both the raw low-dose input and the baseline CycleGAN model. These Quantitative results show that the RDBCycle-GAN outperforms the vaniella cycleGAN in terms of all metrics. Its outputs are much closer to the full-dose reference than the vaniella cycleGAN and quarter-dose inputs.

**TABLE I.** The table lists the imaging parameters for each patient.

| Patient ID | Number of Slices | Size of FOV | KVP | Exposure Time (ms) | X-ray Tube Current (mA) |
| --- | --- | --- | --- | --- | --- |
| L067 | 412 | 370 | 100 | 500 | 234.1 |
| L096 | 518 | 430 | 120 | 500 | 327.6 |
| L109 | 318 | 400 | 100 | 500 | 322.3 |
| L143 | 422 | 440 | 120 | 500 | 416.9 |
| L192 | 428 | 380 | 100 | 500 | 431.6 |
| L286 | 398 | 380 | 120 | 500 | 328.9 |
| L291 | 531 | 380 | 120 | 500 | 322.7 |
| L310 | 402 | 380 | 120 | 500 | 300.0 |
| L333 | 432 | 400 | 100 | 500 | 348.7 |
| L506 | 399 | 380 | 100 | 500 | 277.7 |

**TABLE II.** Quantitative Comparison OF LDCT Input, Vanilla CycleGAN, AND THE Proposed RDBCycleGAN-CBAM.

| Parameter | LDCT |  |  | Vanilla cycleGAN |  |  | RDB cycleGAN |  |  |
| --- | --- | --- | --- | --- | --- | --- | --- | --- | --- |
| | Mean ± Std | Median | IQR | Mean ± Std | Median | IQR | Mean ± Std | Median | IQR |
| PSNR | $34.170 \pm 4.586$ | 33.146 | 2.932 | $35.8 \pm 4.7$ | 35.1 | 3.100 | $38.143 \pm 4.544$ | 37.108 | 2.885 |
| SSIM | $0.893 \pm 0.060$ | 0.897 | 0.049 | $0.918 \pm 0.045$ | 0.921 | 0.036 | $0.946 \pm 0.033$ | 0.950 | 0.026 |
| RMSE | $63.760 \pm 14.131$ | 58.954 | 11.244 | $49.8 \pm 10.9$ | 46.7 | 8.900 | $40.288 \pm 8.583$ | 37.244 | 6.825 |
| MAE | $45.398 \pm 10.814$ | 41.331 | 8.576 | $36.9 \pm 8.2$ | 34.1 | 6.600 | $30.027 \pm 6.846$ | 27.363 | 5.370 |
| NMSE | $0.008 \pm 0.005$ | 0.006 | 0.003 | $0.005 \pm 0.003$ | 0.004 | 0.002 | $0.003 \pm 0.002$ | 0.002 | 0.001 |
| SNR | $21.625 \pm 2.087$ | 22.177 | 1.925 | $23.6 \pm 2.1$ | 24.100 | 1.900 | $25.599 \pm 2.022$ | 26.180 | 1.895 |
| Edge Error | $(4.24 \pm 0.99) \times 10^7$ | $3.89 \times 10^7$ | $7.55 \times 10^6$ | $(3.10 \pm 0.72) \times 10^7$ | $2.95 \times 10^7$ | $5.90 \times 10^6$ | $(2.41 \pm 0.49) \times 10^7$ | $2.24 \times 10^7$ | $4.05 \times 10^6$ |
| EKI | — | — | — | $0.392 \pm 0.025$ | 0.390 | 0.028 | $0.428 \pm 0.020$ | 0.427 | 0.024 |

### B. Bland-Altman Analysis

A Bland–Altman analysis on the N = 421 paired test images for RDBCycleGAN-CBAM shows a clear positive bias in both metrics: for PSNR the mean difference (bias) is MDPSNR ≈ +3.974 dB with 95% limits of agreement [+3.786, +4.162] dB, and for SSIM the bias is MDSSIM ≈ +0.053 with 95% LoA ≈ [−0.001, +0.108]. The PSNR LoA are extremely narrow (span ≈ 0.38 dB), indicating a highly consistent ≈ 4 dB improvement for virtually every image, while the SSIM LoA are wider, indicating more image-dependent variability in structural gains (most images improved, a few showed negligible change). Importantly, no case exhibited meaningful degradation (the lower LoA are essentially zero)

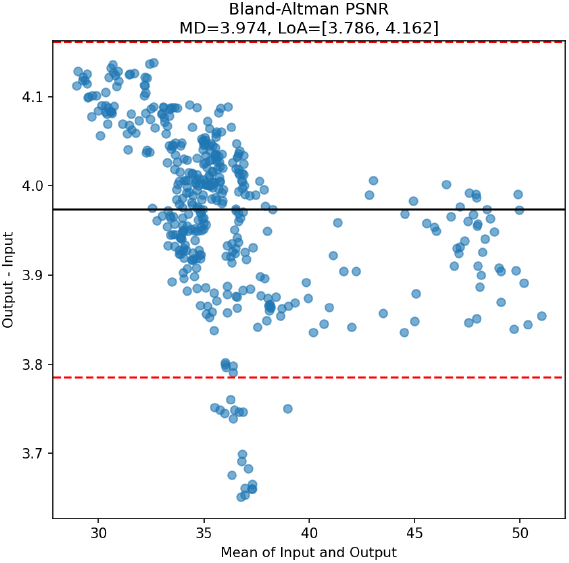
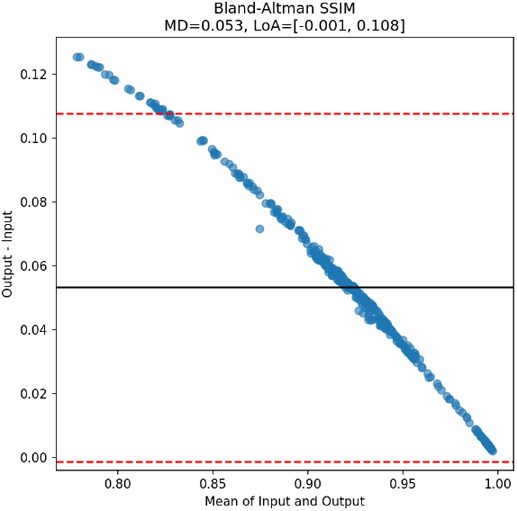

### C. Paired-Lines Analysis

The paired-line plots for N = 421 test images show a uniformly positive effect of the denoising method: every paired line slopes upward for both PSNR and SSIM (421/421 improved), indicating no cases of no-change or degradation.

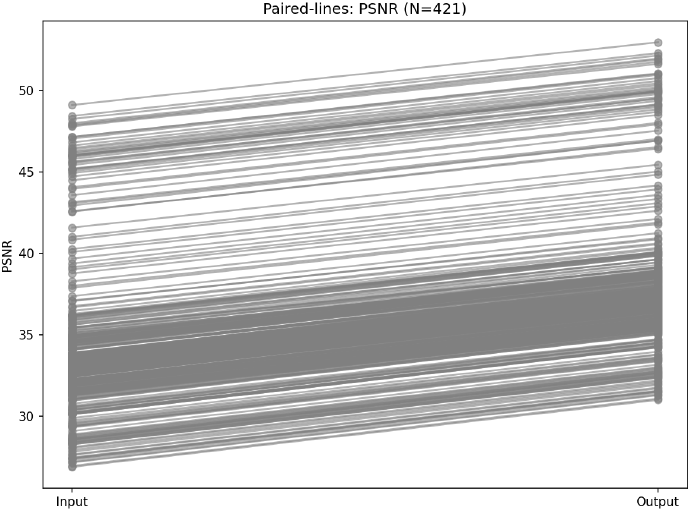
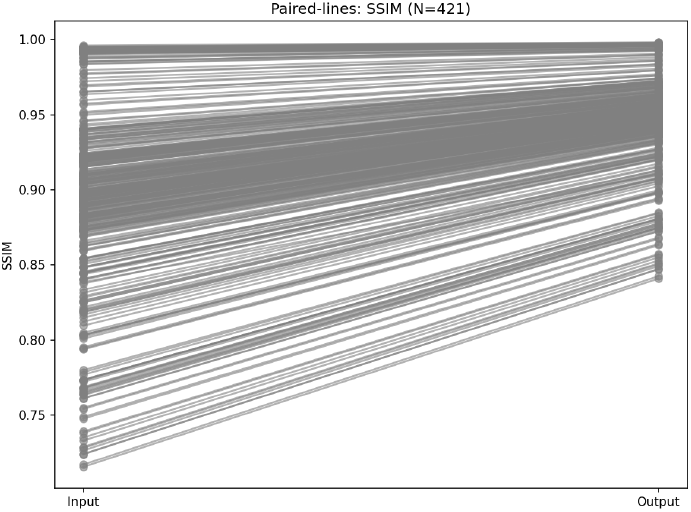

### D. PSNR and SSIM Distribution Analysis

The boxplot (PSNR) and violin plot (SSIM) analyses demonstrate a clear and consistent improvement in image quality from input to output images. The PSNR distribution is entirely shifted upward for the output images, while the SSIM violin becomes markedly narrower and concentrated near higher values, signifying both improved quality and reduced variability.

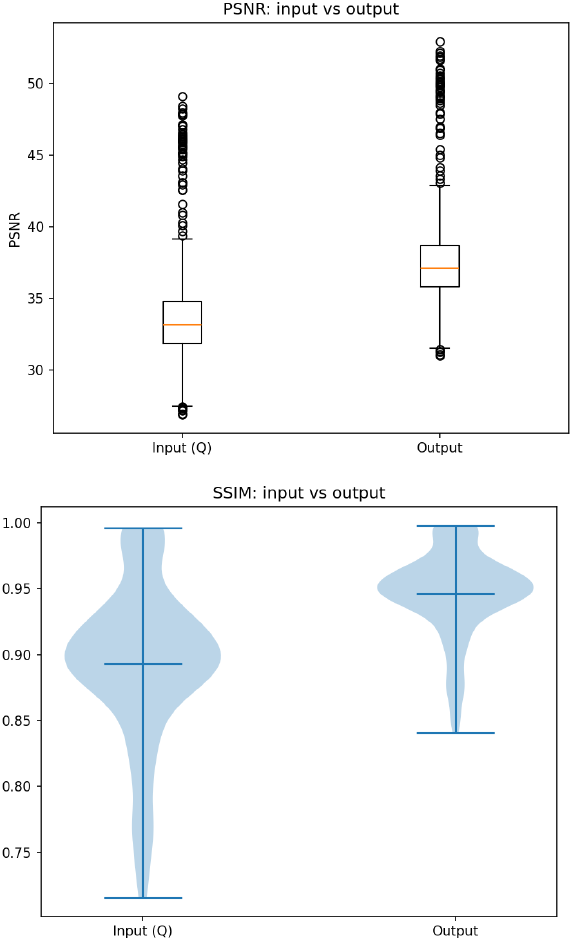

### E. Qualitative Evaluation

The qualitative comparison in Fig.8. demonstrates the effectiveness of the proposed CycleGAN-based denoising approach in improving low-dose CT image quality. Compared with the quarter-dose inputs (left column), the CycleGAN-denoised images (right column) exhibit substantial noise reduction and improved visual clarity, while closely matching the anatomical appearance of the full-dose reference images. Quantitatively, the quarter-dose inputs achieve PSNR values of 36.41, 32.97 and 31.26 dB with corresponding SSIM values of 0.91, 0.86 and 0.84 when compared to the full-dose images, Table III. After denoising, the CycleGAN outputs show consistent improvements, the results showed in Table III corresponds to an average PSNR gain of approximately 4 dB across all slices, indicating effective and uniform noise suppression. The SSIM improvements, which range from moderate to substantial depending on the slice, indicate enhanced preservation of structural information and anatomical details. Overall, both visual inspection and quantitative metrics confirm that the proposed method successfully reduces noise while maintaining structural fidelity, producing denoised images that are substantially closer to the full-dose reference than the original quarter-dose inputs.

**Fig. 8.**
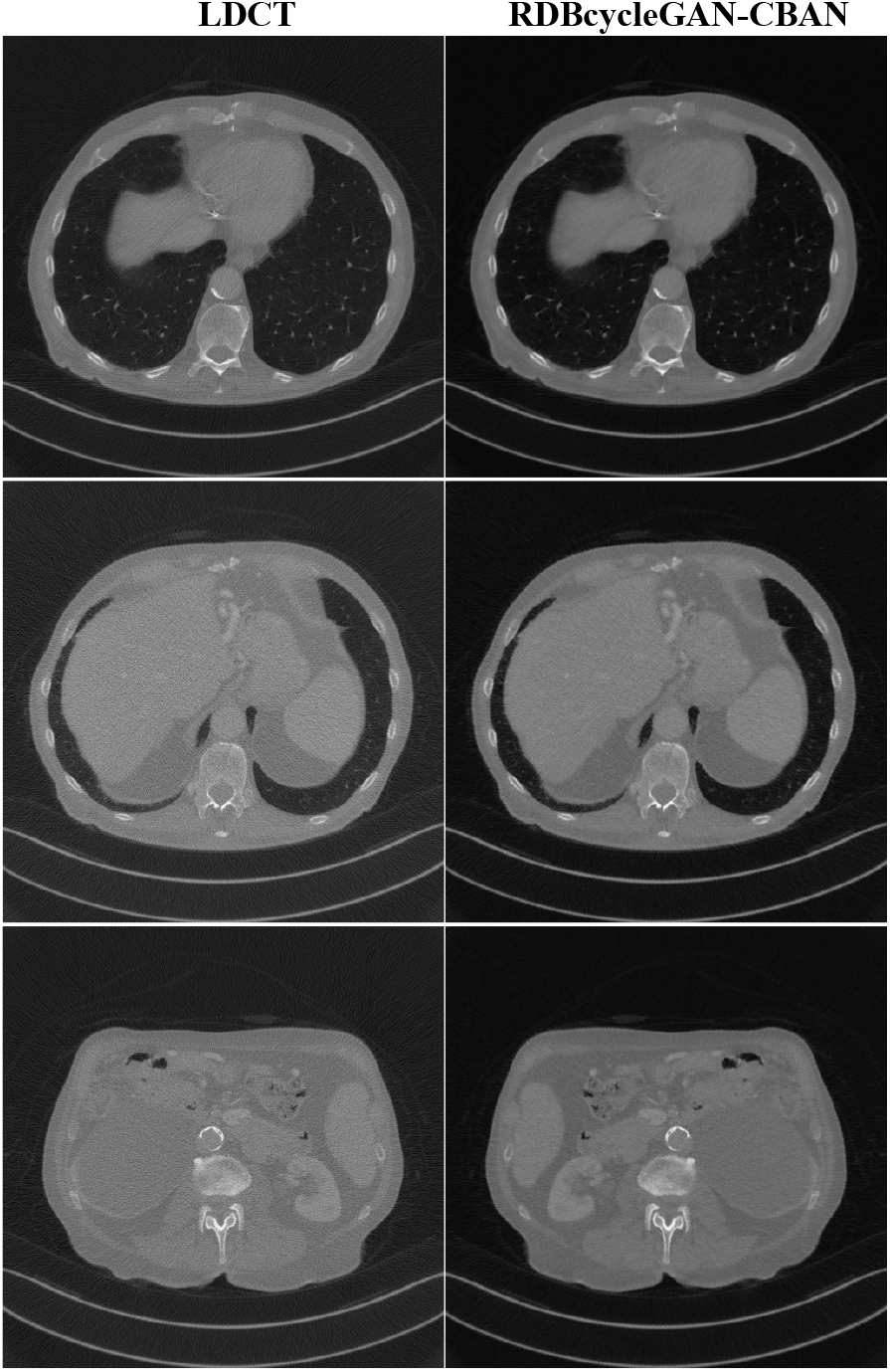
Qualitative comparison of abdominal CT denoising results.

**TABLE III.** Quantitative Results OF THE Full Images&ROIs Corresponding TO Fig.8.

| Slice | LDCT |  | RDB-CycleGAN-CBAM |  |
| --- | --- | --- | --- | --- |
|  | PSNR (dB) | SSIM | PSNR (dB) | SSIM |
| 1 | 36.41 | 0.91 | 40.35 | 0.95 |
| 2 | 32.97 | 0.86 | 36.94 | 0.90 |
| 3 | 31.26 | 0.84 | 35.29 | 0.92 |

As indicated by the red circles in Fig. 9, the quarter-dose image exhibits pronounced noise and grainy texture that obscure fine anatomical details. In contrast, the RDBCycleGAN-CBAM output demonstrates substantial noise suppression and improved tissue homogeneity while preserving structural features and lesion boundaries, closely resembling the full-dose reference, see Fig. 11. These visual results illustrate the model’s ability to reduce low-dose noise without introducing noticeable blurring or artificial structures.

**Fig. 9.**
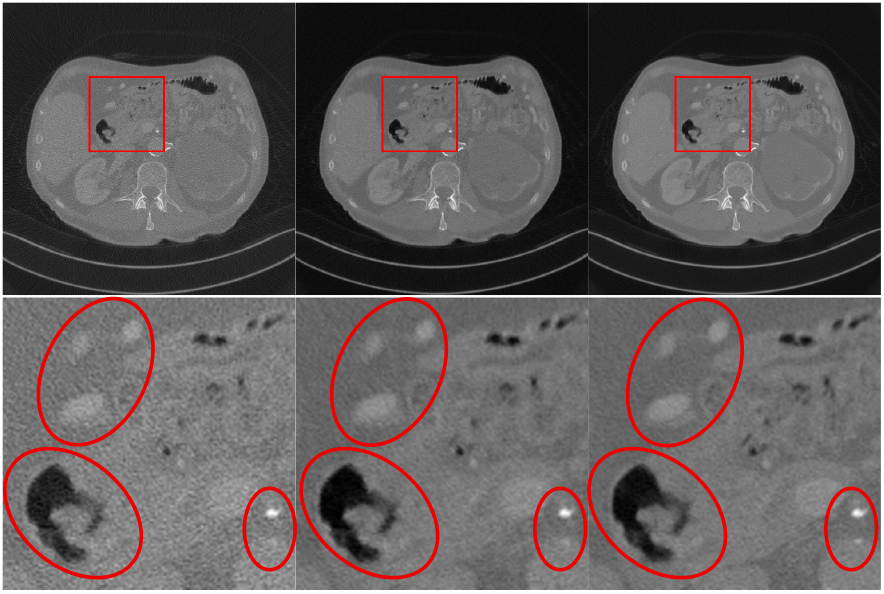
Qualitative comparison of abdominal CT denoising results with ROIs.

Visual inspection confirms substantial noise suppression while preserving anatomical structures such as vessel boundaries and organ interfaces. Residual maps in Fig.10 indicate removal of high-frequency noise without large-scale structural distortion.

**Fig. 10.**
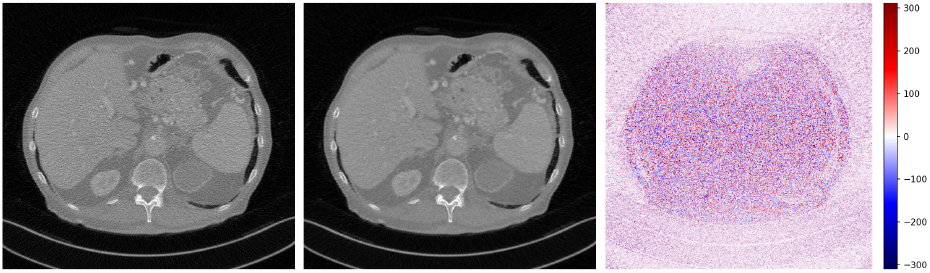
Example axial slice illustrating residual map results

### A. Statical Evaluation

To assess statistical significance of the improvements in PSNR and SSIM, we first tested whether the paired differences (output minus input) were normally distributed. Shapiroa-Wilk tests[26] on the paired differences (output minus input) indicate a significant departure from normality for both PSNR (W = 0.9519, p = 1.79 × 10^−10^) and SSIM (W = 0.9515, p = 1.57 × 10^−10^); accordingly, nonparametric inference was performed using the Wilcoxon signed-rank test[27]. For both metrics the Wilcoxon test statistic is 0 with p ≈ 1.01 × 10^−70^, and the rank-biserial effect size is 1.00[28], which together indicate that every single paired comparison favored the CycleGAN output (no ties or negative differences). The mean paired improvements are large and precise: PSNR increased by +3.9737 dB (95% CI [3.9646, 3.9826] dB) and SSIM increased by +0.05319 (95% CI [0.05060, 0.05579]). In practical terms, these results in Table IV show an overwhelmingly consistent and statistically decisive improvement in both pixel-wise fidelity (PSNR) and structural similarity (SSIM) for the CycleGAN-processed images relative to the quarter-dose inputs.

**TABLE IV.**
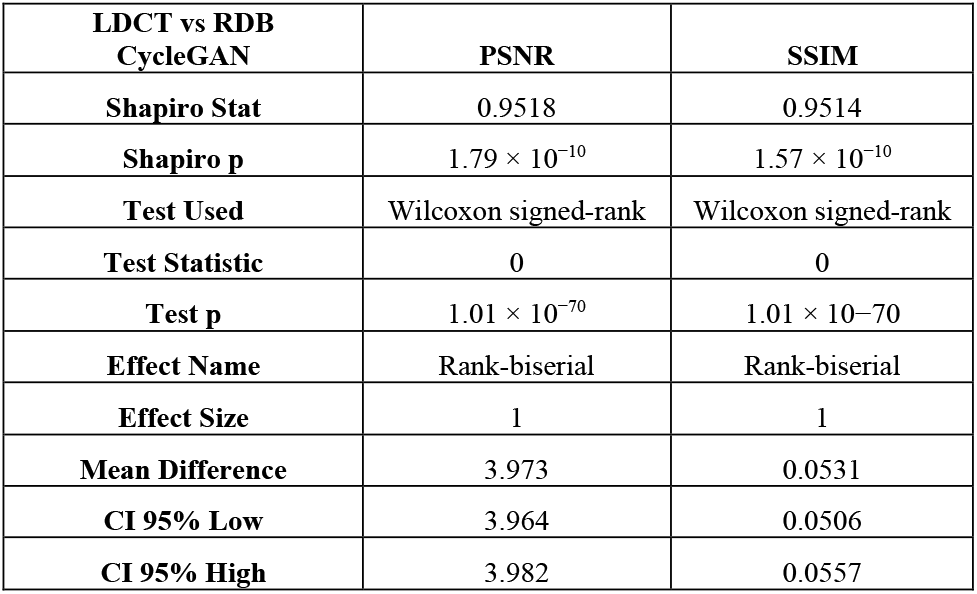
Summary OF Statical Evaluation Results.

| LDCT vs RDB<br>CycleGAN | PSNR | SSIM |
| --- | --- | --- |
| Shapiro Stat | 0.9518 | 0.9514 |
| Shapiro p | $1.79 \times 10^{-10}$ | $1.57 \times 10^{-10}$ |
| Test Used | Wilcoxon signed-rank | Wilcoxon signed-rank |
| Test Statistic | 0 | 0 |
| Test p | $1.01 \times 10^{-70}$ | $1.01 \times 10^{-70}$ |
| Effect Name | Rank-biserial | Rank-biserial |
| Effect Size | 1 | 1 |
| Mean Difference | 3.973 | 0.0531 |
| CI 95% Low | 3.964 | 0.0506 |
| CI 95% High | 3.982 | 0.0557 |

### B. Quantitative Comparison With Other Methods

The denoising gains achieved by our RDBCycleGAN-CBAM are comparable to or better than those reported for recent methods on the AAPM Mayo dataset. In our experiments RDBCycleGAN-CBAM increased mean PSNR by ≈ +3.97 dB and SSIM by ≈ +0.053. By comparison in Table V, the literature reports more modest improvements. For example, Zhou et al. [29] report that RED-CNN, CycleGAN and IdentityGAN each boost PSNR by roughly +2–3 dB with SSIM gains of ∼ 0.03– 0.04, and that GAN-CIRCLE yields ≈ +3.0 dB PSNR and +0.038 SSIM. Similarly, Wolterink et al. [30] found that a WGAN-VGG model gave about +0.81dB PSNR and +0.034 SSIM relative to LDCT. Chen et al. [31] report their SKFCycleGAN achieving +2.3 dB PSNR and 0.047 SSIM on the Mayo LDCT data. In summary, RDBCycleGAN-CBAM’s nearly +4 dB PSNR gain is on par with the best prior models, and its +0.053 SSIM increase is larger than those previously reported—indicating very competitive denoising performance, especially in preserving image structure.

**Table V.**
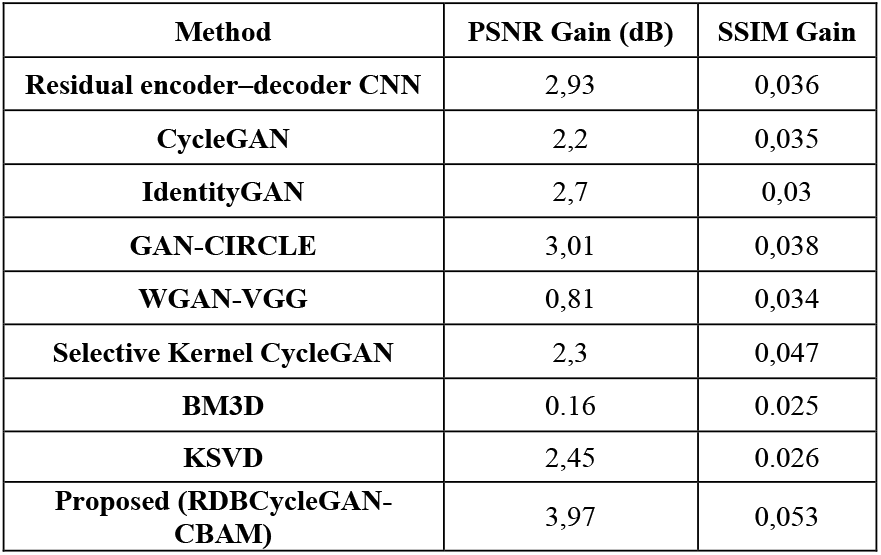
Quantitative Comparison With Other Methods.

| Method | PSNR Gain (dB) | SSIM Gain |
| --- | --- | --- |
| Residual encoder-decoder CNN | 2,93 | 0,036 |
| CycleGAN | 2,2 | 0,035 |
| IdentityGAN | 2,7 | 0,03 |
| GAN-CIRCLE | 3,01 | 0,038 |
| WGAN-VGG | 0,81 | 0,034 |
| Selective Kernel CycleGAN | 2,3 | 0,047 |
| BM3D | 0.16 | 0.025 |
| KSVD | 2,45 | 0.026 |
| Proposed (RDBCycleGAN-CBAM) | 3,97 | 0,053 |

**TABLE VI.**
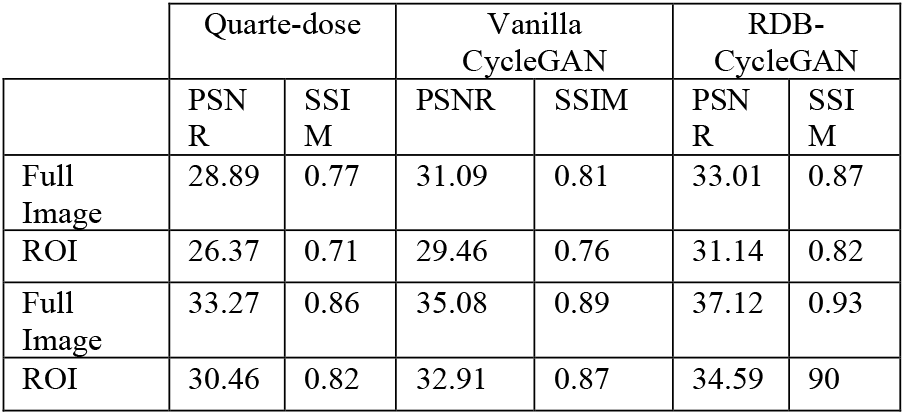
Quantitative Results OF THE Full Images&ROIs Corresponding TO Fig. 11.

## V. DISCUSSION

RDBCycleGAN-CBAM integrates several complementary modules that improve LDCT denoising. Residual learning at multiple scales (ResNet blocks, RDBs, and a global residual) stabilizes training and directs the network to model noise/artifact corrections rather than reconstruct the entire image. Additionally, dilated convolutions enlarge the effective receptive field, so the model uses larger anatomical context to distinguish noise from real structure. Furthermore, dense/RDB connections promote feature reuse and improved information flow in the bottleneck, helping preserve subtle details. Finally, CBAM attention adaptively focuses capacity on diagnostically important channels and spatial regions (edges, vessels).

On standard LDCT benchmarks (NIH-AAPM Mayo dataset), RDBCycleGAN-CBAM attains PSNR/SSIM on par with or exceeding recent methods in Table V, while qualitatively preserving sharper edges and finer details relative to many supervised and unpaired baselines. As shown in Fig. 11 The proposed method demonstrates improved noise reduction and edge preservation, resulting in images that are much closer to routine-dose quality. Compared with supervised CNNs that often oversmooth, our adversarially trained model better balances perceptual detail and pixelwise fidelity; perceptually oriented methods (e.g., WGAN-VGG) may trade PSNR for visual quality, whereas the inclusion of RDBs and CBAM here improves both objective and perceptual metrics.

**Fig. 11.**
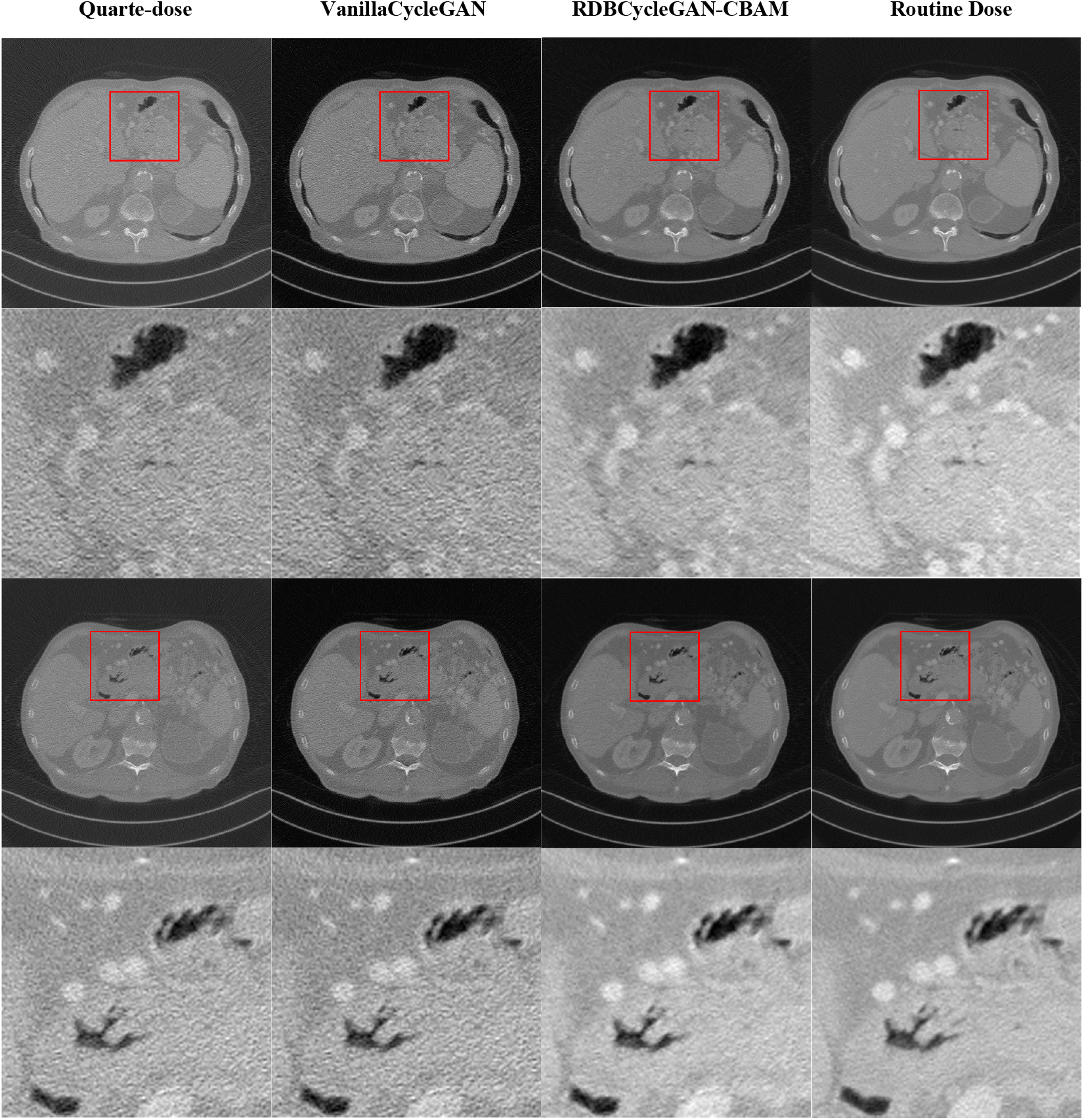
Qualitative comparison of LDCT denoising. From left to right: quarter-dose, Vanilla CycleGAN, RDBCycleGAN-CBAM (proposed), and routine-dose reference. Red boxes indicate ROIs; corresponding zoomed views are shown below. The proposed method achieves stronger noise suppression and better edge preservation, yielding images closer to routine-dose quality.

GAN-based denoisers carry a nonzero risk of hallucinating plausible but incorrect structures, particularly in very low-signal regions; we observed occasional, spatially localized spurious textures in a small subset of cases. There is also an inherent smoothing vs. detail trade-off: pixel losses favor PSNR while adversarial/perceptual losses favor sharper appearance. Model generalization depends on data diversity domain shifts (scanner/protocol/pathology) may degrade performance so extensive, multi-center validation is required before clinical use.

Stable training requires careful loss-weighting and monitoring; we used Adam for ∼80–100 epochs and found mixed-precision training helpful. The model is large (tens of millions of parameters across two generators) and benefits from high-memory GPUs (e.g., V100/A100) for efficient training; inference on modern GPUs is fast enough for near-real-time slice-wise processing. By improving LDCT image quality, RDBCycleGAN-CBAM has potential to support dose reduction, but clinical deployment requires rigorous reader studies and prospective validation to rule out subtle biases or rare hallucinations. With appropriate validation and regulatory review, the method could contribute to safer, higher-quality CT imaging in practice.

## VI. CONCLUSION

We introduce RDBCycleGAN-CBAM, a cycle-consistent GAN that integrates residual-dense blocks (RDBs), dilated convolutions, and a CBAM attention module to improve feature extraction and structural preservation in LDCT denoising. We developed a reproducible training pipeline (PyTorch, data augmentation, mixed precision) and trained the model on quarter-dose ↔full-dose 2D slices, yielding stable convergence with cycle and adversarial losses. On 421 test slices, the model increased PSNR by ≈ +3.97 dB and SSIM by ≈ +0.053 versus quarter-dose inputs, with corresponding reductions in RMSE/MAE/NMSE and improved edge preservation (higher EKI). Improvements were statistically significant (Wilcoxon test, p < 0.05). The model is efficient at inference (slice-wise processing on modern GPUs), making near-real-time deployment feasible, and its compact design supports practical integration into radiology pipelines.

Integrating RDBs and attention into a CycleGAN framework yields images that are quantitatively closer to full-dose references and that retain fine anatomical detail, see Fig. 11. The observed ∼4 dB PSNR gain, and consistent Wilcoxon results indicate a uniform improvement across the test set, which is important for clinical reliability. Edge-preserving metrics corroborate the qualitative observation that the method reduces noise without excessive blurring.

By algorithmically enhancing quarter-dose CT images to near full-dose quality, this approach could enable substantial radiation reductions (e.g., 75% dose reduction) in appropriate clinical contexts, facilitating safer repeat imaging and broader screening while maintaining diagnostic utility. However, this potential depends on rigorous clinical validation.

Limitations include a residual risk of GAN-induced hallucinations in very low-signal regions, sensitivity to domain shifts (scanner/protocol/pathology), and substantial training compute requirements.

Future work will focus on: (1) volumetric and inter-slice models to improve 3D consistency; (2) multi-center reader studies and task-based evaluations (lesion detectability); (3) robustness across dose regimes and scanners via domain adaptation; and (4) architectural and deployment optimizations (projection-domain integration, diffusion/transformer alternatives, compression for edge inference).

RDBCycleGAN-CBAM offers a practical, vendor-neutral post-processing route to improve LDCT quality while preserving diagnostic detail. With further validation and regulatory review, it has the potential to contribute meaningfully to dose-reduction strategies in clinical CT imaging.

---

**Osman Assaf**, Results-oriented Biomedical Engineer holding a Bachelor’s, and a Master’s degree in Biomedical Engineering (Medical Imaging specialization) from Boğaziçi University (Robert College). Completed MRI modality training at the UIH HQ Training Center in Shanghai, combining strong hands-on modality expertise with over four years of academic and scientific research on MRI and CT systems. Demonstrated a proven track record in the installation, commissioning, and start-up of multiple uMR and uCT imaging systems. Experienced in resting-state fMRI (rs-fMRI) processing and Default Mode Network (DMN) identification.

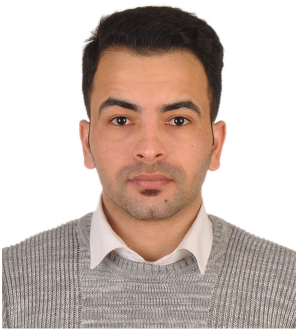

Prof. Dr. **Albert Güveniş** received his PhD. degree from Drexel University, Philadelphia, USA. He then worked as a researcher and a faculty member at the Radiology Dept. in the Hospital of the University of Pennsylvania, Philadelphia, USA in the area of Positron Emission Tomography. He then joined Boğaziçi University.

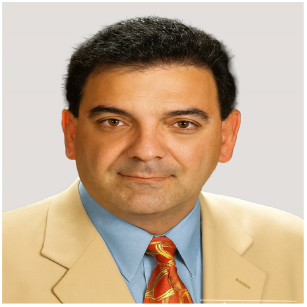

